# A feedback loop able to enlarge the brain for 2.4 myr without Darwin’s selective survival

**DOI:** 10.1101/053827

**Authors:** William H. Calvin

**Affiliations:** Department of Psychiatry and Behavioral Sciences, University of Washington, 725 9th Avenue, Box 2605, Seattle WA 98104-2086 USA, and the UCSD/Salk Center for Academic Research and Training in Anthropogeny, La Jolla CA 92037 USA.

## Abstract

The rapid three-fold enlargement of the hominin brain^1,2^ began about 2.3 million years ago (myr) as Africa dried and grass replaced brush, creating great savannas^3^. Seeking an amplifying feedback loop, I analyzed the lightning-brush-fire ecology for grazing animals in a grassy burn scar^4^. Discovering the new grass by exploring brush byways could promote a population boom–but only after grass-specialized herbivores evolved from mixed feeders^5^ at 2.4 myr. When the brush returned several decades later, the grazer boom would turn to bust, squeezing numerous descendants back into the core grasslands. Meat-eating *Homo* species would boom and bust when grazers did, enriching the core in whatever alleles were earlier concentrated in the brush fringe catchment zone for that boom. This return migration for *Homo* is what creates the amplifying feedback loop that speeds brain enlargement rate, likely up to the mutation rate limit. It also promotes trait hitchhiking: any brush-relevant allele, not just those for hunting, can experience amplifying feedback merely by hanging out in the catchment zone^4^. The shade offered by brush would have been the default location for cooperative nurseries, time-consuming food preparation, and toolmaking. Increased behavioral versatility correlates with larger brain size and the more versatile brains of a current generation need only spend more-than-average time in the boom’s catchment zone for this recursive evolutionary process to keep average brain size increasing via assortative mating. This helps account for the time when enlargement began, why it was linear, when it ended, and why it slowed in Neanderthals and in Asian *Homo erectus*. Without utilizing Darwin’s selective survival, the feedback loop makes advance room for “free” future functionality in the cerebral cortex, likely relevant to the evolutionary emergence of our structured intellectual functions^6^ such as syntax, contingent planning, games, and logic.

---

How do the early stages of a new neocortical function—say, syntax—evolve via the selective survival of useful mutations, the best-known aspect of Darwin’s natural selection? A bump-by-bump cortical enlargement, where each bump comes from a chance enlargement of an underlying functional map that is conserved by selective survival, is what the standard evolutionary argument expects. But it need not be so specific: comparative mammalian studies^7^ suggest that the easy path to a local enlargement is an overall neocortical increase, resulting in extra space for other functions that have no immediate payoff, e.g., if better visual acuity is conserved by selective survival, a side effect might be improved auditory discrimination. Here I analyze a second nonspecific mechanism, an allele-amplifying feedback loop also generating free space; it can operate quickly and runs well without utilizing selective survival.

The scatter plot of hominin brain size over time (Fig. 1a) shows four major features: a long period of near-stasis in the Pliocene where brain size in the Australopithecines remained in the range for the extant great apes; a fast 3.3x rise during the Pleistocene; an apparent doubling of growth rate during the most recent 0.2 myr; and as part of a more general gracilization with agriculture, a sudden 10–17% reduction during the Holocene^8^, the equivalent of more than 0.3 myr down the trend. However, the gracilization suggests a relaxation of maintaining selection for robustness, not retrogression of new function.

**Fig. 1.**
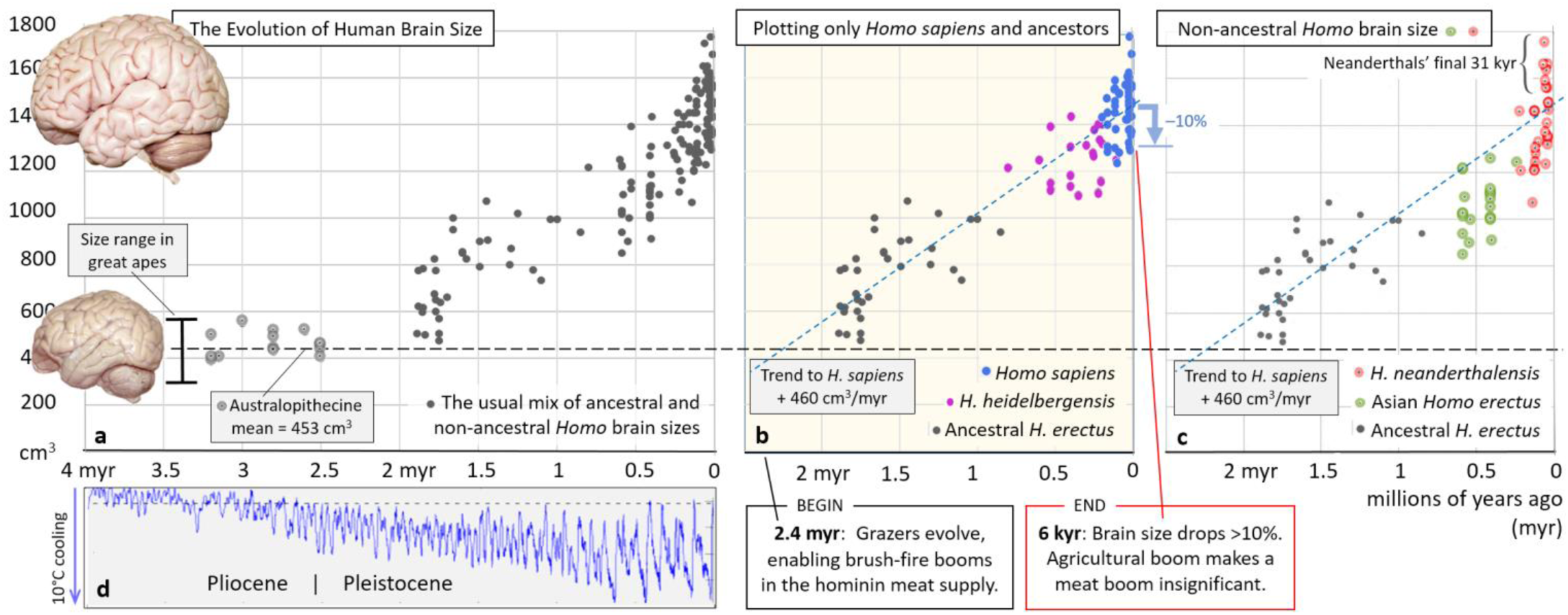
The evolution of human brain size. **a.** Australopithecines had brains little larger than those of the great apes (a bonobo brain is illustrated, to the same scale as the human brain). After a long period of stasis, hominin brain size^2^ (N=175) increased 3.3 times, with the enlargement rate appearing to double in the last 0.2 myr. **b**. However, this piecewise linearity disappears if one excludes known non-ancestors. The average Australopithecine brain size (N=14) is shown by the dashed line at 453 cm^3^. The dashed blue line (repeated in **c**) intercepting it at 2.3 myr is extrapolated from the least-squares fit to ancestral endocranial capacity (N=125) between 12 kyr and 1.9 myr, yielding a trend of 460 cm^3^/myr. The blue down-arrow shows the drop in size^8^ during the Holocene. **c.** Plotting the omitted brains shows that the non-ancestors (N=36) lag behind the ancestral trend line, explaining the piecewise linearity seen in **a**; the Neanderthals have a substantial brain enlargement only in their final 31 kyr, perhaps from the interbreeding^11^ with *H. sapiens*. **d.** Proxy of ocean temperature^12^ shows ice age fluctuations in climate, averaged over 57 sediment-core sites.

Since *Homo sapiens* appears^9^ at 0.2 myr, the late doubling is of particular interest. However, Fig. 1a includes Neandertals and Asian *Homo erectus*; they distort the scatter plot on which previous analyzes of piecewise enlargement have relied^2^. In Fig. 1b, these known non-ancestors are omitted. Ancestral brain enlargement appears quite regular at 460 cm^3^ every million years, using a least-squares fit from 12 kyr back to 1.9 myr. There no longer appears to be a late spurt for *H. sapiens*; rather the question becomes one of why the rate remained constant.

The line fit in Fig. 1b intercepts the endocranial capacity axis at 1491 cm^3^. Even allowing for the looser fit of brain to bone in modern *H. sapiens*, this seems 10% high when compared to modern brain size^1^ averaging 1330 cm^3^. However, the Fig. 1 data set^2^ ends at 12 kyr; after then (blue down arrow in Fig. 1b), human cranial capacity decreased enough during the agricultural Holocene^6^ to make the match reasonable.

At the other end, the fitted line extrapolates back to Australopithecine brain sizes at 2.3 myr. Grazers evolved from mixed-feeders^5^ at 2.4 myr; this allows brush-fire booms (Fig. 2) in the hominin meat supply to commence. Grazers’ population size is limited by encroaching brush (neither browsers nor mixed feeders should boom when some brush becomes temporary grassland), so grazer booms become possible when new grasslands appear after a brush fire, and thus meat booms in *Homo*.

**Fig. 2.**
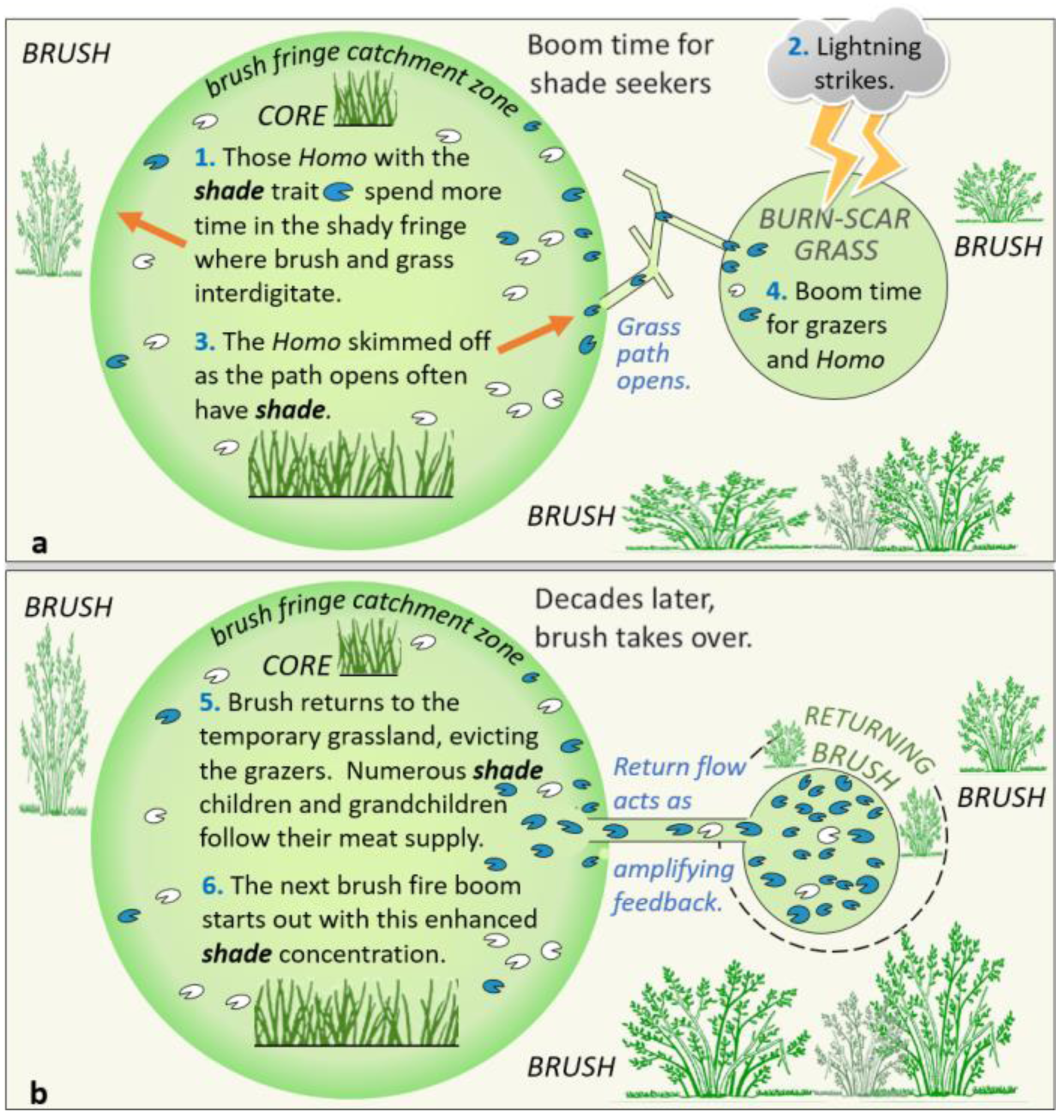
The boom-and-bust feedback loop promotes hitchhiking by traits such as shade-seeking. **a**. After grazers experience a population boom upon discovering the burn scar’s temporary grassland, the meat-eating hominin population will secondarily experience a boom. **b**. Alleles with a concentration gradient between the core and the brush fringe catchment zone, such as those utilizing shade, will boom and then enrich the core with brush-relevant alleles decades later via the return flow. This creates amplifying feedback.

The non-ancestors are plotted separately in Fig. 1c, which shows that Asian *H. erectus* caused the sag in the middle of Fig. 1a. Slowing of enlargement is also seen in the Neanderthals if one ignores the six largest Neanderthal brains, which are all from the final period between 71 kyr and the 40 kyr disappearance^10^. This Neanderthal spurt, perhaps four times faster than the *H. sapiens* trend, suggests a different enlargement regime in that final period, such as interbreeding 11 with *H. sapiens*.

Contrary to a century of expectations, there are no obvious changes in enlargement rate that might mark the successively colder depths of the glacial periods (Fig, 1d) during the last million years^12^, or the major advances in stone toolmaking at million year intervals, or the appearance of new *Homo* species at 0.8 and 0.2 myr.

Such prediction failures would cast doubt on many of the previous explanations offered for brain enlargement but for the following: looking at evolutionary development as a complex system suggests we are simply seeing a rate-limited process, as when the speed of an assembly line is limited by the slowest supplier of parts. Suppliers of other parts might have fluctuations in delivery rate but one would not observe this in the output rate unless that rate became slower than that of the current rate-limiter.

In this manner, changing selection pressures for brain size might not show up in the enlargement record if such a ramped “glass ceiling” was limiting the rate. Amplifying feedback always has such limiters. While there is no need for a limiting rate to remain constant, the one for brain enlargement appears to have done so, and in spite of every terrestrial environment on earth undergoing major change (Fig. 1d) time and again during the Pleistocene. While we await even more fossils for mid-range Fig. 1b, should we be looking off-earth for the rate that does not adapt?

If all of the relevant brain-sizing genes had become fixed, as are most genes, there would have been little raw material for a Darwinian competition, not until mutation made another allele. The simplest way to tweak a gene’s normal function is via a single nucleotide polymorphism (SNP), created when a passing neutron of cosmic origin knocks out a single base pair in the germ line and gene repair then makes a mistake. Genome wide, there are about 60 new SNPs in each newborn human, reflecting gamete mutations during its parents’ lives^13^. Should the modified DNA triplet code for a different amino acid (thus creating an allele), the resulting protein may fold differently and so modify the function served, such as more/less, sooner/later during development.

With so many neutron sources in deep space, intervals between cosmic ray SNPs will be random but the mean arrival rate should not change on the million-year time scale. We must assume that what began about 2.4 myr were brain enlargement processes that are inherently faster than the mutation rate. Amplification feedback can be very fast until pinned by a limiter (what allows binary computation).

Two rate-limiter candidates have been identified in human evolution: the brain’s share of the blood supply had to keep increasing at the expense of something else, likely prolonged digestion^14^, and increased head size^15^ had to be accommodated by shortening gestation and/or widening the birth canal bottleneck. However, producing new alleles is the ultimate rate limiter for brain enlargement; there may well have been situations where there could not be sudden spurts as new adaptations proved their worth, given that the feedback loop was already pinning enlargement rate to this limit.

As background for the *Homo* boom-and-bust feedback loop (Fig. 2), note that most species vastly over-reproduce; their surplus-to-replacements offspring usually die before reproductive maturity unless a boom occurs. Thus most selective survival operates not on the adult but on the more numerous immature; juvenile success determines which survive to reproduce at all.

But natural selection does not operate solely by selective survival; it can also operate by selective reproductive success, as in those species variants where twinning rate temporarily increases with diet improvements^16^ at the end of a drought. Selective range expansion is another way to achieve a similar effect.

Brush fires are far more frequent than droughts or other climate change events; lightning strikes that occur in the dry season can create large burn scars that promptly grow grass. If this auxiliary grassland opens up for the grazers in the core’s brush fringe, most of their surplus-to-replacement offspring can survive to reproduce. Then comes the bust as the brush returns. A selective expansion coupled with return migration (Fig. 2) can produce a substantial shift in core allele proportions (“gene frequency”) in a few generations^4^, what would take many generations of selective survival operating on a peripheral fraction of the population.

By hanging out in the catchment zone for the boom of a major food source, hitchhiking is enabled for hominin traits that tend to concentrate there, often for reasons other than predation. For example, shade-seeking in brush could serve to concentrate traits for toolmaking, food preparation, and the communal rearing aspect of eusociality^4^. The boom and its feedback loop would then amplify those traits in the core, once the bust occurred. Because of repeated amplification, even weak traits can be quickly enhanced.

If behavioral versatility concentrates in the catchment zone, this amplifying feedback loop would have affected brain enlargement. For example, those versatile enough to skillfully hunt both grass-eating grazers and leaf-eating browsers would spend more time in the brush, making them more likely to be there when a gateway opened to the hidden grassland. Those specializing in open-terrain confrontational scavenging^17^, or running an herbivore to exhaustion^18^, would often miss the opportunity.

Because brain size is strongly heritable^19^, bootstrapping to a new normal via assortative mating can be done by the more versatile hanging out in the catchment zone for the boom time feedback loop. An evolutionary arms race is not needed and there is no need to keep proving the worth of a slightly bigger brain via selective survival. While we normally speak of selection “pressure,” this aspect of natural selection seems more permissive, better able to account for long-run advantages.

Building up behavioral versatility over the last 2.4 myr includes improving the hand-eye coordination for the “get set” ballistic movements^20^, such as the increasingly delicate hammering needed for the fine serrated cutting edges of Achulean-style tools^21^ and for accurate throwing from ever greater distances. Early *Homo erectus/ergaster* appeared on the East African scene by 1.9 myr with a shoulder already adapted for throwing projectiles^22^.

As there are no second chances (if a projectile misses its target, dinner runs away), and because many targets are not at a standard distance, planning well during “get set” to throw accurately on the first try is a demanding task for the brain^6^, much more so than targeting the future position of a moving target. Furthermore, herds will move back sooner when hunters again approach, requiring improvements in throwing accuracy just to maintain the hit rate at the new move-back distance, encouraging practice sessions. The law of large numbers means that devoting more neurons to the timing of projectile release is one way to reduce timing jitter, improving accuracy^20^. The genes improving throwing were exposed to selective survival when there was a meat-rich diet but may have also benefitted from both types of nonspecific neocortical enlargement.

Such elaborate (and often novel to the instance) muscle command sequences could benefit from a motor equivalent of working memory^23^, ideally a flexible workspace in the cerebral cortex for the detailed “get set” planning of the movements that are too quick for delayed sensory corrections to guide them. Throwing, hammering, clubbing, kicking, and speech might utilize this ballistic workspace.

With time, a new innate movement specialization–say, hammering–might take root in the flexible workspace, perhaps at the expense of losing some space for planning throws, thereby decreasing throwing accuracy. But since there is a considerable spread in brain size in a given generation, those individuals with a brain sufficiently larger than the current average will possess the original amount of flexible workspace despite their space commitment to the new hammering specialization. Larger-than-average also serves as a preadaptation, making it easier to map yet another neocortical innovation in those individuals of the current generation with sufficiently larger brains.

As with chemistry’s autocatalytic processes, this allele amplification loop has failure modes. Had grazers been hunted to extinction, browsers and mixed-feeders would still have provided an adequate meat supply for *Homo*, new alleles would have continued to form, but there would have been no boom to run the allele amplification loop.

Loop failures may have influenced non-ancestral *Homo* (Fig. 1c) as they emigrated into environments where the feedback loop rarely operated. The Rift path at 1.8 myr from equatorial Africa to the 42°N plate boundary at Dmanisi^24^ would have allowed the feedback loops along the way to repeatedly enrich emigrants with brush-relevant alleles. But subsequently moving into forested areas would leave the feedback loop behind. Very large grasslands, such as the Steppes of Central Asia, may not have enough adjacent brush to create significant booms after lightning strikes, slowing allele shifts in their core. Rift valleys, and other river valleys with grassy hillsides and brush at the broad bottom, should allow the temporary grassland population to be a more significant proportion of the combined population.

A role for amplifying feedback suggests some new answers to the traditional questions about the tempo and mode of evolution, such as switching on brain enlargement and providing the setup for the two great Out of Africa expansions. While variation and selection will happen elsewhere as well, that along the Rift valley has the amplification advantage, providing a fast track able to preempt many late-arriving adaptations from elsewhere.

While the concepts of versatility and intelligence overlap, note that what the loop amplifies may simply be the correlates of shade-seeking in migratory bands, not specifically the tasks that dominate modern IQ assessment^25^ such as the span of working memory, quick decision-making of high quality, abstraction, and analogy.

Trait hitchhiking for versatility does open up a new way of thinking about an evolutionary extravagance, such as the large gap in intellect between great apes and preagricultural humans that so puzzled Alfred Russel Wallace^26,27^ in 1869 when selective survival seemed to be the only evolutionary tool. Structured cognitive functions are extravagant by great ape standards and it is difficult to make selective survival arguments for their early phases.

Where female mate choice is possible, male reputation becomes of particular interest because exogamy results in an uninformed adolescent female. Facial expression, gaze, pantomime, gesture, and short-sentence protolanguage can all be used to gossip about “Who did what to whom, where, when, and with what means?” But this is laborious when adult hands are busy with shade tasks. A speedier verbal version of gossip^28^ and other long-sentence utterance needs local structuring conventions^29^ (grammar and syntax) if a listener is to quickly understand the longer complex strings such as “I think I saw him leave to go hunting” with its four nested verbs.

Any verbal structured version of gossip would thus be routinely overheard in the shade by infants and emulated by young children, much as in the modern developmental sequence for language^4,30^. While there is little in this protolanguage-to-language example that is exposed to selective survival, the shady setting provides preferred access to selective expansion’s boom-and-bust loop that runs independent of selective survival.

The free space aspect might be said to create opportunities to randomly self-organize novel representations–maps in search of a function–perhaps yielding a novel cognitive capability in those with a larger-than-average brain. Then, with time and further enlargement, the new ability would become available to the entire population: this is bootstrapping across generations as the larger-brain, more versatile adults provide structured examples for all to emulate during the sensitive periods for soft wiring in childhood.

To summarize the feedback loop itself, there is the need for specialized grazers in order to produce a boom after a brush fire, a need for some isolation spanning decades, and a return path when the brush takes over again. For trait hitchhiking in the feedback loop, it needs co-located heritable traits in the brush’s fringe with grassland such as toolmaking, food preparation, and the communal nursing of infants^4^. This clustering permits assortative mating of the more versatile to increase mean brain size, though rate-limited by mutation rate.

This establishes a rationale for when enlargement began (grazers evolve), why it was linear (steady accumulation of favorable alleles, fixed by amplification as fast as the SNP clock ticks), when it ended (agriculture enlarges core population to such an extent that meat booms become insignificant), and why enlargement slowed in side branches (compared to the African Rift, there are fewer settings able to amplify a new allele with a meat boom).

Trait hitchhiking now seems promising to explore as an alternative evolutionary path for syntax and the other structured intellectual functions^6^: contingent planning, chains of logic, games with rules constraining moves, analogies that extend to parables, polyphonic music, and creativity’s eureka moments when incoherent mental assemblies become coherent fits, good to go.

For exploring an evolutionary extravaganza, examining settings may be more productive than the usual focus on increments in usefulness. As sexual selection did for the extravagant peacock tail, feedback loops can surprise us with progressions that keep going automatically–as in that afore-mentioned neocortical preadaptation for the next new thing, in all the children who are above average.

## Acknowledgements

I thank my La Jolla colleagues for two decades of expert discussions about human evolution, with long-term CARTA funding from the Mathers Foundation. Human and bonobo scaled photographs thanks to T.W. Deacon. I thank my University of Washington colleagues James J. Anderson, Katherine Graubard, and Charles D. Laird for discussions on the manuscript.

**Supplemental data:** Spreadsheet for Fig. 1 is available on-line at Nature [link].

## References

1. Holloway, R.S. The evolution of the hominid brain. In Henke, W., & Tattersall, I. (eds.), Handbook of Paleoanthropology, pp.1961–1987. Berlin Heidelberg: Springer-Verlag (2015) | doi:10.1007/978–3–642–39979–4_81.

2. Shultz, S., Nelson, E., & Dunbar, R.I.M. Hominin cognitive evolution: identifying patterns and processes in the fossil and archaeological record. Phil Trans Roy Soc B 367: 2130–2140 (2012) | doi:10.1098/rstb.2012.0115.

3. deMenocal, P.B. African climate change and faunal evolution during the Pliocene-Pleistocene. Earth and Planetary Science Letters 220: 3–24 (2004).

4. Calvin, W.H. Eusociality and other improbable evolutionary outcomes can be accelerated by trait hitchhiking in boom-bust feedback loops. Preprint at bioRxiv (2016) | doi:10.1011/053819.

5. Cerling, T.E. et al. Dietary changes of large herbivores in the Turkana Basin, Kenya from 4 to 1 Ma. Proc Natl Acad Sci U S A. 112: 11467–11472 (2015) | doi:10.1073/pnas.1513075112

6. Calvin, W.H. A Brief History of the Mind. New York: Oxford University Press (2004).

7. Finlay, B.L., & Darlington, R.B. Linked regularities in the development and evolution of mammalian brains. Science 268: 1578–84 (1995).

8. Henneberg, M., & Steyn, M. Trends in cranial capacity and cranial index in Subsaharan Africa during the Holocene. American Journal of Human Biology 4: 473–479 Fig. 4. (1993) | doi:10.1002/ajhb.1310050411.

9. McDougall, I., Brown, F.H., & Fleagle, J.G. Sapropels and the age of hominins Omo I and II, Kibish, Ethiopia. Journal of Human Evolution 55, 409–420 (2008) | doi:10.1016/j.jhevol.2008.05.012

10. Higham, T. et al. The timing and spatiotemporal patterning of Neanderthal disappearance. Nature, 512: 306–309 (2014) | doi:10.1038/nature13621.

11. Pääbo, S. Neanderthal Man: In Search of Lost Genomes. New York: Basic Books (2014).

12. Lisiecki, L.E., & Raymo, M.E. A. Pliocene-Pleistocene stack of 57 globally distributed benthic δ180 records. Paleoceanography 20: PA1003 (2005) | doi:10.1029/2004PA001071.

13. Kong, A., et al. Rate of de novo mutations and the importance of father’s age to disease risk. Nature 488, 471–475 (2012) | doi:10.1038/nature11396.

14. Aiello, L.C., & Wheeler, P. The expensive-tissue hypothesis: the brain and the digestive system in human and primate evolution. Current Anthropology 36: 199–221(1995) | Link: www.jstor.org/stable/2744104.

15. Rosenberg, K. R. (1992), The evolution of modern human childbirth. Am. J. Phys. Anthropol., 35: 89–124 (1992) | doi:10.1002/ajpa.1330350605.

16. Heape, W. Note on the fertility of different breeds of sheep, with remarks on the prevalence of abortion and barrenness therein. Proc. Roy. Soc. Lond. 65, 99–111 (1899).

17. Bunn, H.T. Hunting, power scavenging, and butchering by Hadza foragers and by Plio-Pleistocene Homo. In: Stanford, C.B., & Bunn, H.T., editors. Meat-eating and Human Evolution. New York: Oxford University Press, pp. 199–218 (2001).

18. Lieberman, D.E., Bramble, D.M., Raichlen, R.A., & Shea, J., J. Brains, brawn, and the evolution of human endurance running capabilities. In Grine, F.E., Fleagle, J.G., & Leakey, R.E., editors. The First Humans – Origin and Early Evolution of the Genus Homo. Amsterdam: Springer Netherlands; pp. 77–92 (2009).

19. Witelson, S.F., Beresh, H., & Kigar, D.L. Intelligence and brain size in 100 postmortem brains: sex, lateralization and age factors. Brain 129: 386–398 (2006) | doi:10.1093/brain/awh696.

20. Calvin, W.H. A stone’s throw and its launch window: timing precision and its implications for language and hominid brains. J. Theor. Biol. 104: 121–135 (1983).

21. Dibble, H.L., et al. Major fallacies surrounding stone artifacts and assemblages. J Archaeol Method Theory (2016) | doi:10.1007/s10816–016–9297–8.

22. Roach, N.T., Venkadesan, M., Rainbow, M.J., & Lieberman, D.E. Elastic energy storage in the shoulder and the evolution of high-speed throwing in Homo. Nature 498: 483–486 (2013) |doi:10.1038/nature12267.

23. Gallivan, J.P., Johnsrude, I.S., & Flanagan, J.R. Planning ahead: object-directed sequential actions decoded from human frontoparietal and occipitotemporal networks. Cerebral Cortex (2015) | doi:10.1093/cercor/bhu302.

24. Messager, E., Lordkipanidze, D., Kvavadze, E., Ferring, C.R., & Voinchet, P. Palaeoenvironmental reconstruction of Dmanisi site (Georgia) based on palaeobotanical data. Quaternary International 223: 20–27 (2010) | doi:10.1016/j. quaint.2009.12.016.

25. Calvin, W.H. Abrupt climate jumps and the evolution of higher intellectual functions during the Ice Ages. In Sternberg, R.J., ed., The Evolution of Intelligence. Erlbaum, pp. 97–115 (1999).

26. Wallace, A.R. The Malay Archipelago: the land of the orang-utan and the bird of paradise; a narrative of travel, with studies of man and nature. London: Macmillan (1869), p.204.

27. Bickerton, D. More than Nature Needs: Language, Mind, and Evolution. Cambridge MA: Harvard University Press (2014).

28. Dunbar, R.I.M. Gossip in evolutionary perspective. Review of general psychology 8: 100–110 (2004) | doi:10.1037/1089–2680.8.2.100

29. Calvin, W.H., & Bickerton, D. Lingua ex Machina. Cambridge MA: MIT Press (2000), p.83.

30. Kuhl, P.K. Early language acquisition: cracking the speech code. Nature Reviews Neuroscience 5, 831–843 (2004) | doi:10.1038/nrn1533

